# Nanoscale membrane curvature sorts lipid phases and alters lipid diffusion

**DOI:** 10.1101/2020.09.23.310086

**Authors:** Xinxin Woodward, Matti Javanainen, Balázs Fábián, Christopher V. Kelly

## Abstract

The precise spatiotemporal control of nanoscale membrane shape and composition is the result of complex interplay of individual and collective molecular behaviors. Here, we employed single-molecule localization microscopy and computational simulations to observe single-lipid diffusion and sorting in model membranes with varying compositions, phase, temperature, and curvature. Supported lipid bilayers were created over 50-nm radius nanoparticles to mimic the size of naturally occurring membrane buds, such as endocytic pits and the formation of viral envelopes. The curved membranes recruited liquid-disordered lipid phases while altering the diffusion and sorting of tracer lipids. Disorder-preferring fluorescent lipids sorted to and experienced faster diffusion on the nanoscale curvature only when embedded in a membrane capable of sustaining lipid phase separation at low temperatures. The curvature-induced sorting and faster diffusion even occurred when the sample temperature was above the miscibility temperature of the planar membrane, implying that the nanoscale curvature could induce phase separation in otherwise homogeneous membranes. Further confirmation and understanding of these results are provided by continuum and coarse-grained molecular dynamics simulations with explicit and spontaneous curvature-phase coupling, respectively. The curvature-induced membrane compositional heterogeneity and altered dynamics were achieved only with a coupling of the curvature with a lipid phase separation. These cross-validating results demonstrate the complex interplay of lipid phases, molecular diffusion, and nanoscale membrane curvature that are critical for membrane functionality.

**Significance:** Nanoscopic membrane organization and dynamics are critical for cellular function but challenging to experimentally measure. This work brings together super-resolution optical methods with multiscale computational approaches to reveal the interplay between curvature, composition, phase, and diffusion in model membranes. We report that curvature can induce phase separation in otherwise homogeneous membranes and that the phase-curvature coupling has a direct implication on lipid mobility. This discovery advances our understanding of the fundamental membrane biophysics that regulate membrane activities such as endocytosis and viral budding.

## Introduction

Cell plasma membranes contain thousands of distinct lipid species with spatial heterogeneity in composition and shape. Natural membrane compositions near a phase coexistence manifest functional membrane domains that are more and less structured and resemble liquid ordered (L_o_) and liquid disordered (L_d_) phases [31, 35, 36, 42]. Membrane domains are critical for cell functions such as protein sorting, cell signaling, membrane budding, and retrovirus replication through the regulation of local variations in the membrane composition [10, 17, 35, 37, 47]. Caveolae, for example, are cholesterol and sphingolipid-rich endocytic buds that have diverse roles in membrane compositional regulation, trafficking, force sensing, and endocytosis on the sub-100-nm length scale [26, 29, 30]. The coupling of membrane phases and curvature on live cells remains unconfirmed yet likely crucial for diverse cellular functions.

Model membranes of known lipid mixtures provide a means to reveal how the membrane composition affects function. Coexisting L_o_ and L_d_ phases can be created by combining a sterol, a phospholipid with a high chain-melting temperature, and a phospholipid with a low chain-melting temperature below the mixture’s miscibility transition temperature (*T*_mix_) [43, 45]. Phase-separated model membranes have shown curvature-dependent phase separations in tubules [16, 32, 38, 41], small vesicles [21], and supported lipid bilayers (SLBs) [27, 28, 52]. In select cases, the membrane shape has been shown to induce a phase separation that was not observed on planar membranes [32, 38]. In model membranes, the region of greater curvature (*i*.*e*., smaller radius) concentrates disordered lipids as the L_d_ phase has a lower bending rigidity than the L_o_ phase [2]. Interestingly, however, endocytic processes and viral budding in live cells are typically correlated with L_o_-preferring lipids, such as cholesterol and sphingomyelin [46], which may occur if the area of the ordered phase is significantly smaller than the disordered phase or if the order-preferring lipids and proteins have a greater spontaneous curvature [24]. Typically, single lipids are too small relative to the membrane curvature for the curvature to significantly alter their distribution in the membrane without the collective action of lipid phases [5, 7, 21, 22]. The physical principles that govern the sorting of lipids to *<*100-nm membrane buds remains a mystery fundamental to both cellular homeostasis and pathophysiology. This manuscript reports the temperature-dependent diffusion and sorting on membranes to better understand the interplay of lipid mobility, phase separation, and curvature.

Lipid diffusion in the L_d_ phase is up to 10*×* faster than lipids in the L_o_ phase, depending on the membrane composition, the fluorescent lipids tracked, and the substrate topography [6, 8, 11, 13, 33, 50]. For example, DPPE-Texas Red in SLBs composed of DiPhyPC, DPPC, and cholesterol at molar ratios of 2:2:1 had a diffusion coefficient (1.8 *±* 0.5)*×* greater in the L_d_ phase than the L_o_ phase [48]. The diffusion in the two phases became indistinguishable upon increasing the temperature or cholesterol content, as is associated with shorter tie-lines separating the phases. For example, SLBs composed of DiPhyPC, DPPC, and cholesterol at molar ratios of 1:1:2 demonstrated lipid phase separation, as evident by the partitioning of DPPE-Texas Red, but no difference in the DPPE-Texas Red diffusion was observed between the two phases [48].

Experimental limitations and the underlying topological manifold result in the motion of particles on curved membranes appearing non-Brownian [14]. Restrictions on the observable lag times and localizations precision result in the measured diffusion becoming increasingly non-local, in the sense that observed particle motion averages patches of the membrane. Consequently, the lipid mobility becomes reflective of the average membrane environment with a loss of nanoscale information. Nonetheless, modern single-particle localization and computational methods are converging to reveal spatially varying membrane behaviors at physiological length scales.

The influence of membrane curvature on single-lipid diffusion has been studied in well-defined geometries. Diffusion typically slows on cylindrical membranes when the radii are smaller than 50 nm [9]. But the effects of curvature on lipid diffusion are correlated to the fluorophore location on the labeled lipid; head-group-labeled lipids diffuse 3*×* faster on flat versus curved membranes, whereas tail-labeled lipids diffuse at the same rate on flat and curved membranes [49]. Recent experimental and computational capabilities to measure diffusion and sorting at the nanoscale are beginning to reveal an intertwined network of contributing factors affecting membrane phenomena, including the effects of hydrodynamics, geometry, projection, lipid packing, and measurement time. For example, membrane curvature affects the thickness, area-per-lipid, and the order parameter of the lipid hydrocarbon chains [51].

In this manuscript, the diffusion and sorting of lipids relative to nanoscale membrane curvature and phase separation are quantified with experimental and computational methods. Membrane curvature was experimentally generated by creating phase-separated SLBs over 50-nm radius nanoparticles on planar microscopy coverslips (Fig. 1). The concentration of DPPE-Texas Red on the curved membrane was measured *via* diffraction-limited images and analyzed for lipid sorting. Single-molecule localization microscopy and single-particle tracking (SPT) were performed to track individual DPPE-Texas Red molecules and to measure their curvature- and phase-dependent mobility. The disorder-preferring fluorescent lipids were more concentrated on the curved membrane than the surrounding planar bilayer when embedded in a membrane capable of sustaining lipid phase separation. This sorting occurred regardless of the phase of the immediately surrounding bilayer, but sorting was more pronounced if the membrane bud was immediately surrounded by an L_o_ phase. The single-lipid diffusion was up to 2.2*×* faster on curved versus planar membranes, with a dependence on the length of the phase-separating tie-lines. These experimental results were verified in computational simulations of phase separation, including continuum and molecular dynamics simulations with explicit or spontaneous phase-curvature coupling, respectively. The sorting of disordered lipids to the curvature and the effects of the curvature propagating onto the surrounding planar membrane were consistently observed. These diverse approaches independently measure sorting and diffusion to demonstrated the lipid phase coupling to nanoscale membrane bending.

**Figure 1:**
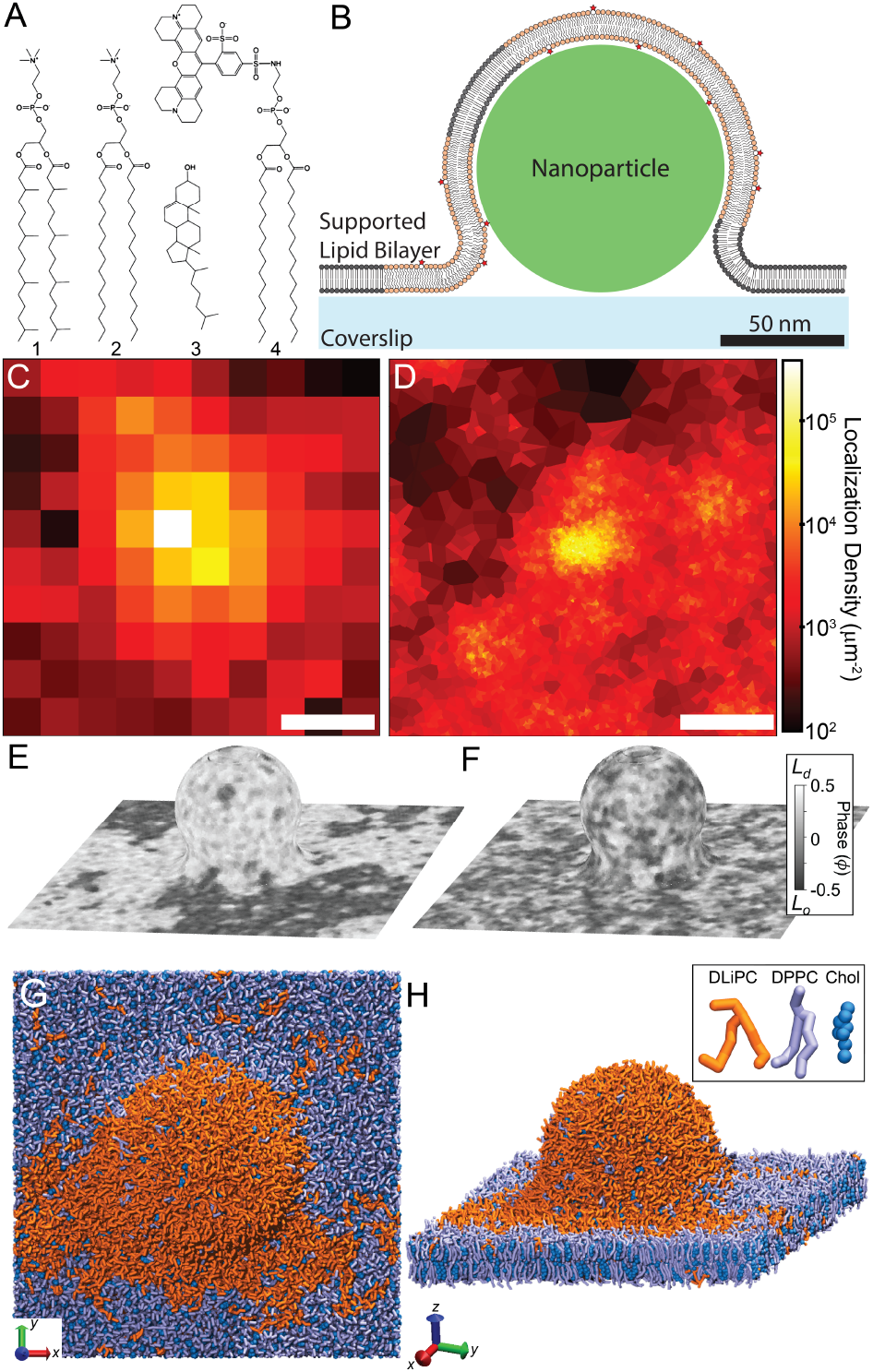
Curvature in phase-separated lipid bilayers was studied both experimentally and computationally. (A) The lipids used include (1) DiPhyPC, (2) DPPE, (3) cholesterol, and (4) DPPE-Texas Red. (B) The lipid phases and single-lipid diffusion was experimentally measured relative to the engineered, nanoparticle-induced membrane curvature. (C) Diffraction-limited and (D) super-resolution microscopy of SLBs with engineered curvature revealed the sorting and diffusion of lipids; scale bars represent 200 nm. A continuum simulation modeled the curved, phase-separated membrane at (E) *T* = 15^°^C and (F) *T* = 35^°^C and with explicit phase-curvature coupling, *γ* = 0.1. (G, H) Coarsegrained molecular dynamics simulations spontaneously demonstrated curvature-phase coupling through the lipid sorting and diffusion.

## Results

### Phase separation in planar SLBs

To assess the lipid phases within the membranes, we added fluorescence tracer lipids to ternary mixtures of DiPhyPC, DPPC, and cholesterol. Phase-separated GUVs were burst upon microscopy coverslips to create SLBs patches with distinct lipid phase of similar size as present on the GUVs, *i*.*e*., *>*5 μm wide domains. Bright domains on the membranes indicated the L_d_ phase, dim domains indicated the L_o_ phase, and black regions indicated the lack of an SLB over the coverslip (Fig. S1). Substrate-membrane interactions prevented the diffusion of domains in the SLB and the coalescence of optically resolvable lipid phases in SLBs that were created by small vesicle fusion. The fluorescence contrast between the L_o_ and L_d_ phases reflects the partition coefficient of the fluorescent lipid and the length of the tie-line that separated the two phases. The L_o_ and L_d_ phases were more similar in brightness at higher cholesterol concentrations and at higher temperatures.

The observed *T*_mix_ varies between SLB patches similar to the variation between GUVs in a sample [44]. The *T*_mix_ for planar SLBs with a molar ratio of 1:1:2 DiPhyPC:DPPC:cholesterol was between 28^°^C and 37^°^C, as evident by the stable phase separation at 28^°^C and the phase mixing at 37^°^C (Fig. S2).

The *T*_mix_ for planar 2:2:1 SLBs was greater than 37^°^C, as evident by the consistently sharp phase boundaries. All samples were equilibrated for *>*30 min prior measuring the phase separation or lipid diffusion. The slow large-scale mixing of SLBs resulted in some of the largest domains (*>*10 μm diameter) remaining evident with blurred phase boundaries after equilibration at a temperature greater than *T*_mix_, as described previously [15, 48].

### L_d_-preferring lipids sort to curvature

To measure if nanoscale membrane curvature preferentially recruited lipid phases, we engineered curvature in SLBs by bursting phase-separated GUVs over 50-nm radius nanoparticles and glass coverslips (Fig. 1B). This setup provided a nanoengineered membrane shape while allowing individual lipids to diffuse freely within the SLB. The connectivity of the membrane between the planar SLB and the membrane bud over the nanoparticle was demonstrated both with fluorescence recovery after photobleaching and the presence of uninterrupted of single-lipid trajectories (Fig. S3) [5, 18].

Lipid sorting was induced by membrane curvature with disorder-preferring fluorescent lipids concentrating on the curvature (Fig. 2A). Quasi-single component POPC and DiPhyPC membranes served as a control and normalization for the phase-independent increase in membrane area per pixel, brightness, and single-molecule sorting to curvature (Eq. S4). In all quasi-single component membranes, no apparent sorting of the fluorescent lipids to the curvature was observed; the local increase in membrane brightness at the bud was consistent with an increase in membrane area projected onto the *XY* -imaging plane [49]. In mixed-composition membranes, the lipid phase on the curved membrane (*P*_C_) was compared to the phase of the flat membrane (*P*_F_) immediately surrounding each nanoparticle-supported membrane bud (Eqs. S3 and S4). *P*_C_ and *P*_F_ were measured for varying membrane compositions and temperatures. In SLBs with phase separation, lower temperatures provide a greater range in *P*_F_ values, which is consistent with lower temperatures yielding longer tie-lines and a wider variety in the fluorescent lipid density across the membrane. The sorting of the L_d_ phase to the curved membrane is evident by *P*_C_ being greater than *P*_F_, *i*.*e*. with the disorder-preferring probe localizing at curved regions. The ratio of *P*_C_*/P*_F_ is analogous a curvature-dependent partition coefficient and a previously used quantification of phase sorting [38].

**Figure 2:**
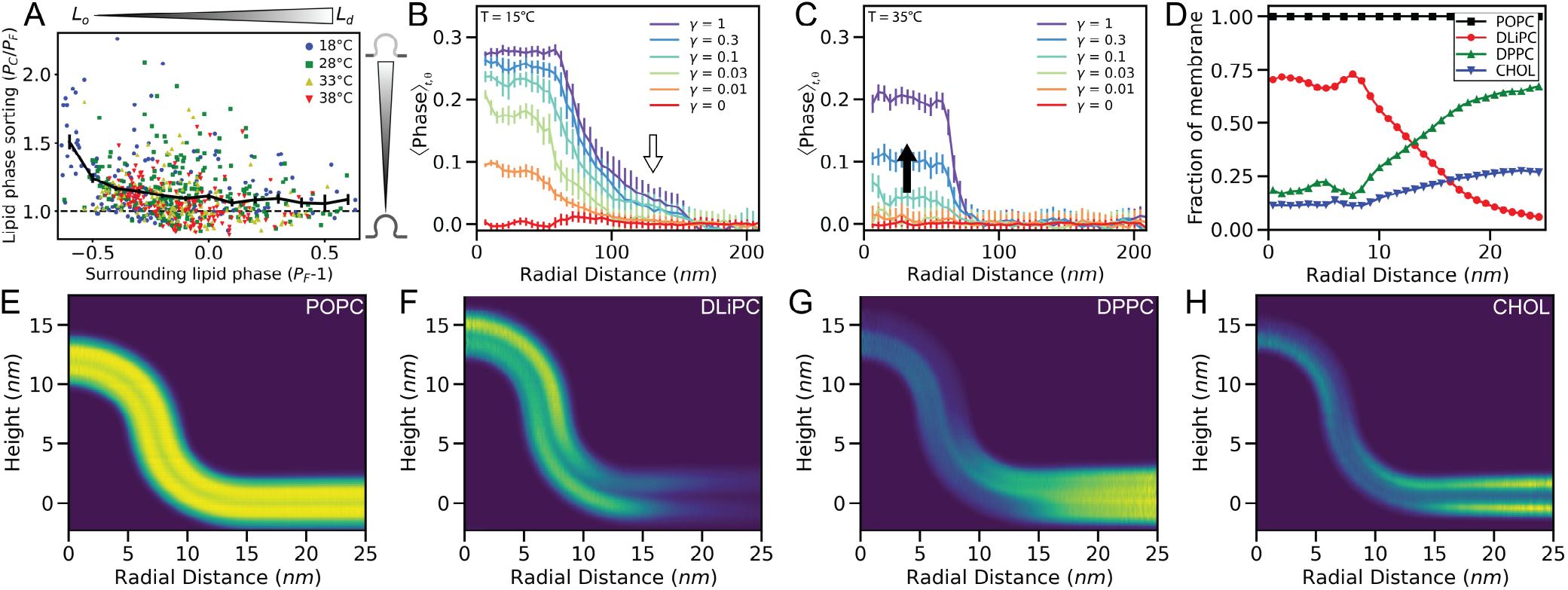
(A) Curvature-induced phase sorting was observed in SLBs with a 1:1:2 molar ratio of DiPhyPC:DPPC:cholesterol. Each data point corresponds to the image of a single membrane bud and the average *P*_C_*/P*_F_ for binned *P*_F_ values are shown in black. Lipids on membranes with spherical, 50-nm radius of curvature are more disordered than the surrounding planar membrane (*i*.*e*., *P*_C_*/P*_F_ *>*1). The curvature-induced phase sorting was greatest when the phase of the flat membrane surrounding the curvature was more ordered (*i*.*e*., (*P*_F_ − 1) *<*0). (B) Continuum simulations of 50-nm radius membrane buds at low temperatures (*T* = 15^°^C) revealed the curvature-preferring L_d_ phase spreading to the surrounding, planar membrane (*white arrow*). (C) Continuum simulations at a high temperatures (*T* = 35^°^C) demonstrated no stable phase separation on the flat membrane, yet concentrated the disordered phase to the curved membrane (*black arrow*). (D–H) Coarse-grained simulations of pure POPC or a mixture of DLiPC, DPPC, and cholesterol revealed the coupling of phase separation and single-lipid diffusion to membrane curvature. (E–H) Normalized 2D histograms show the azimuthally averaged density of lipid beads for each lipid type.

The *P*_C_*/P*_F_ ratio versus *P*_F_ for different phase-separating lipid mixtures and temperatures show no clear change in *P*_C_*/P*_F_ ratio versus temperature (Figs. 2A and S5). The *P*_C_*/P*_F_ ratio was greater than 1 for all temperatures and all values of *P*_F_, which indicates that the curved membrane was consistently more concentrated in the disorder-preferring fluorescent lipid than the surrounding flat membrane. Interestingly, the *P*_C_*/P*_F_ ratio was larger when *P*_F_ was lower, which reveals that the curvature-induced sorting of lipid phases was strongest when the surrounding phase was more ordered.

### Continuum simulation demonstrate curvature-induced lipid domains

To validate the experimental sorting results, we performed computational simulations of a curved membrane with a the local phase governed by the continuum Landau theory and an explicit phase-curvature coupling constant (*γ*) (Eq. S9). The membrane was modeled to have a topography consistent with the experimental, 50-nm radius, nanoparticle-supported SLBs and divided into 3861 distinct cells each with an area of 9 nm^2^ (Figs. 1E, F and S6). Landau phase constants and effective temperatures were chosen to provide two phases with realistic fluctuations in the local composition and phase boundaries, as described in the Supporting Information. Gaussian noise in the phase was added to each cell with each time step to model thermal fluctuations; a larger standard deviation (σ) in this noise represented higher temperature simulations. At low temperature (σ = 0.07), stable domains formed with fluctuations at the phase boundaries (Figs. 1E and 2B). At high temperature (σ = 0.08), rapid fluctuations resulted in no large or stable phases present on the planar membrane (Figs. 1F and 2C). The supplemental material includes movies of these simulations and images of the average phase across the membrane during the simulation with varying temperature and phase-curvature coupling (Movie S1 and Fig. S7). The L_d_ phase was sorted to the curvature in a σ and *γ* dependent manner. At low temperatures, even weak phase-curvature coupling resulted in significant sorting of the L_d_ phase to the membrane bud (Fig. 2B). The phase sorting propagated onto the flat membrane surrounding the bud, which resulted in the planar membrane close to the curvature being more likely to be disordered than the planar membrane far from the curvature (*white arrow*, Fig. 2B). The L_d_ phase was also sorted to the curved membrane even at high temperatures when no large phases were observed on the planar membrane if *γ* ≥ 0.1 (*black arrow*, Fig. 2C).

### Molecular dynamics simulations demonstrate spontaneous phase-curvature coupling

To further validate these results, coarse-grained (CG) molecular dynamics simulations were performed with nanoscale membrane curvature maintained by dummy particles [3]. CG simulations provide single-molecule dynamics from a particular leaflet of the membrane of known shape without perturbations from fluorescent labels. CG simulations included a 5-nm radius hemispherical bud connected to a planar bilayer with a membrane neck of 5-nm minimum radius of curvature (Fig. 1G and H, Movie S2). The membranes had positive Gaussian curvature on the hemispherical bud top, and negative curvature on the neck (Fig. S9A). The simulated lipids diffused and sorted spontaneously relative with to the constant membrane shape according to the Martini force field [25]. The local concentrations of the individual lipid types were azimuthally averaged and either projected onto the *XY* -plane (Fig. 2D), displayed as 2D histograms (Fig. 2E–H), or computed as the real on-surface densities to show the relationship between membrane curvature and leaflet compositions (Fig. S9).

Pure POPC bilayers, which are not capable of inducing a lipid phase separation demonstrated no significant molecular-scale changes in lipid density with curvature (Figs. 2E and S9B). However, for the commonly used phase-separating mixture of DPPC, DLiPC, and cholesterol, the membrane curvature recruited and concentrated the disorder-preferring DLiPC lipids while repelling the order-preferring DPPC and cholesterol. In both leaflets, DLiPC was concentrated on the hemispherical bud and the neck, consistent with experimental and continuum simulation results. The degree of phase separation of the individual leaflets was different, with the upper leaflet inducing a more pronounced sorting (Figs. 2 and S9B).

### Single-lipid diffusion demonstrates the phase-curvature coupling

To test the effects of the curvature-phase coupling on lipid diffusion, experimental and computational SPT was performed to correlate the single-lipid dynamics with the membrane topography. The single-lipid trajectories were experimentally observed in SLBs with engineered membrane buds of 50-nm radius. The lipid phase, curvature, and temperature-dependent diffusion of DPPE-Texas Red was measured (Fig. 3A–D and Table S1).

**Figure 3:**
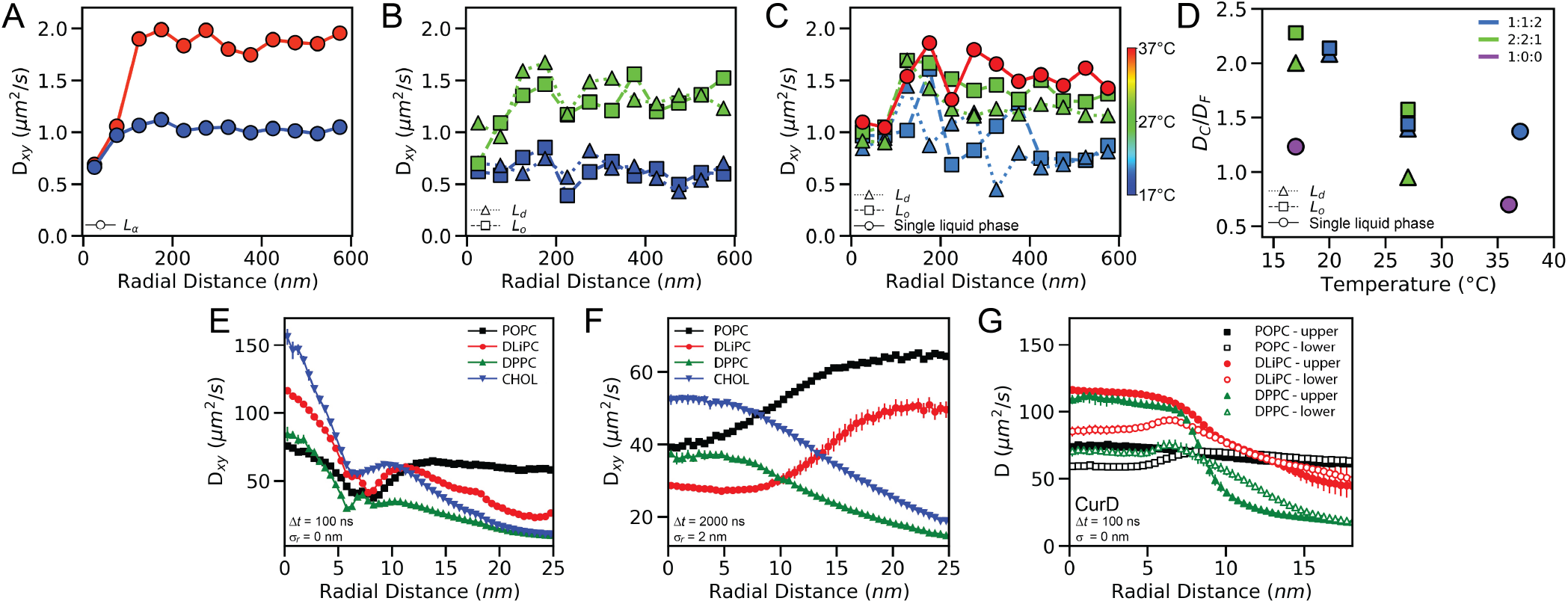
Curvature and phase separation influence the lipid mobility. *D*_*xy*_ as a function of radial distance away from the membrane buds on a planar bilayer was measured for (A) DiPhyPC SLBs and phase-separated SLBs composed of (B) 2:2:1 or (C) 1:1:2 molar ratios of DiPhyPC:DPPC:cholesterol. (D) The diffusion ratio in curved versus flat membranes (*D*_C_*/D*_F_) are shown for varying membrane composition, phase, and temperature. Note that panels (A)–(C) show diffusion projected to the *XY* plane, whereas the data in panel (D) is corrected to account for the real distance along the bud. (A–D) The marker shape indicates the lipid phase. The color indicates the (A–C) the sample temperature or (D) membrane composition. Uncertainty levels for each data point are given in Table S1. The single-lipid step lengths from CG molecular dynamics simulations show (E) diffusion is faster on the curved membrane for all lipid types (Δ*t* = 100 ns; σ_*r*_ = 0), but (F) analysis conditions that scale to the experimental conditions (Δ*t* = 2 μs; σ_*r*_ = 2 nm) incorporate greater spatial averaging coupled with membrane shape and lipid-type distribution differences. (G) Additionally, molecular dynamics results were subjected to a diffusion analysis with geodesic distance-based algorithms (CurD) to reveal the in-membrane diffusion separate from any projection effects.

Diffusion was studied for SLBs of 1:0:0, 1:1:2, and 2:2:1 molar ratios of DiPhyPC:DPPC:cholesterol. Each single-lipid step length through the XY-plane was binned based on the radial lateral distance from the center of the membrane bud (*r*). Rayleigh distribution fitting of the single-step lengths in each bin provided a spatially mapped diffusion coefficient across the topographically varying SLB [18, 19, 49] (Fig. 3, Eq. S9). This SPT analysis method provides greater spatial resolution of lipid diffusion than the conventional fluorescence recovery after photobleaching (FRAP) or mean squared displacement (MSD) versus lag time (Δ*t*) analysis. FRAP typically reports the mean diffusion over a diffraction-limited region of interest, and MSD vs. Δ*t* typically averages together the diffusers’ entire *>*1 μm trajectories (Fig. S8) [20].

The radially dependent diffusion through the *XY* -plane (*D*_*xy*_) varied with membrane topography due to both the curvature-dependent lipid mobility and the geometrical effects of projecting the 3D lipid trajectories into the imaging plane. The contribution due projecting the curved membrane was estimated from Monte Carlo simulations and subtracted to reveal an effective diffusion rate of the lipids on the curved (*D*_C_) and flat membrane (*D*_F_), as described in the Supporting Information S1.10 [49].

As the temperature increased, *D*_F_ increased, as expected [1, 34, 39, 48]. To measure the phase- and curvature-dependent lipid diffusion, membrane buds were grouped based on the phase of the surrounding flat membrane. The individual curvature events were deemed L_d_ or L_o_ phase if *P*_F_ was greater than 1.1 or less than 0.9, respectively. The ratio of *D*_C_*/D*_F_ was consistently higher for systems with longer tie-lines separating the coexisting L_d_ and L_o_ lipid phases (Fig. 3D). The greatest *D*_C_*/D*_F_ ratio was observed in 20^°^C SLBs with a 2:2:1 molar ratio of DiPhyPC:DPPC:cholesterol when the buds were surrounded by L_o_ phase; *D*_C_*/D*_F_ = 2.3 *±* 1.0. In contrast, 25^°^C L_d_ POPC SLBs displayed the smallest *D*_C_*/D*_F_ ratio equal to 0.4 *±* 0.1 (Table S1, Fig. 3D).

All values of *D*_C_*/D*_F_ ratio for ternary mixtures were greater than or equal to one; the lipids in phase-separated membranes diffuse faster if the membrane is curved (Fig. 3D). The *D*_C_*/D*_F_ ratio was larger at the colder temperatures, consistent with the L_d_-preferring lipids sorting to the curvature more when the tie-line separating the phases was longer, as also seen in the continuum models (Figs. 2B and C).

Interestingly, *D*_*xy*_ decreased as *r* increased from 125 nm to 400 nm in ternary mixtures with clear phase separation (*i*.*e*., at lower temperatures and lower cholesterol content) (Fig. 3C). This decreasing *D*_*xy*_ on the flat membrane near the curvature may be due to curvature-seeded L_d_ phase propagation onto the planar SLB surrounding the nanoparticle, as was also seen in both the continuum and CG computational simulations (Fig. 2).

The CG simulations were also analyzed as SPT data, like the experimental data. CG simulations enable analysis with sub-microsecond Δ*t* and with no localization imprecision (σ_*r*_). When analyzed with Δ*t* = 100 ns, the POPC lipids displayed minimal difference in diffusion on the hemispherical bud versus the planar bilayer (*i*.*e*., *r <*3 nm versus *r >*13 nm) (Fig. 3E). The lower *D*_*xy*_ for the POPC lipids at *r* = 8 nm is attributed to the effects of projecting the 3D lipid locations into the *XY* -plane such that motion along the *Z*-axis was incorporated. However, upon the incorporation of a phase-separated lipid mixtures, all lipid types experienced faster diffusion on the membrane bud with Δ*t* = 100 ns, correlated with the curvature-sorted L_d_ phase. Presumably the curvature-associated L_d_ phase provided a lower effective membrane viscosity such that all lipids in the phase-separated membrane diffused faster on the bud.

Analysis of the CG simulations was also performed with conditions mimicking those achieved experimentally (Fig. 3F). Although the observation of spontaneous phase separation in 50-nm buds is not yet computationally feasible, the 5-nm bud simulations were analyzed with mimicking conditions with σ_*r*_ and Δ*t* scaled appropriately. The CG lipid locations at were blurred with σ_*r*_ applied at 10% of the experimental conditions, 2 nm and 20 nm, respectively. The Δ*t* used for the mimicking CG analysis was 1% of that used experimentally, 2 μs and 2 ms, respectively. These analysis conditions resulted in consistent ratios of the localization uncertainty and mean step length to the curvature radius for both the CG simulations and experimental conditions. Additionally, *D*_*xy*_ results presented from CG simulations were corrected with Eq. S6, as done experimentally.

When analyzing the POPC diffusion with longer time steps (Δ*t* = 2 μs) and incorporating a localization uncertainty (σ_*r*_ = 2 nm) to mimic the spatial blurring observed experimentally, *D*_*xy*_ versus *r* for CG POPC closely resembles experimental results for nanoscale buds in quasi-single component supported lipid bilayers (Figs. 3A and F) [18, 49]. With Δ*t* = 2 μs and σ_*r*_ = 2 nm, the diffusion of CG DLiPC resembles that of the CG POPC and experimental DPPE-Texas Red results. But the DPPC and cholesterol diffusion shows significant increase in *D*_*xy*_ on the bud versus the surrounding planar membrane. This result further supports the observation of curvature-phase coupling and motivates the incorporation of a order-preferring fluorescent lipid in future experimental studies.

CG simulations provide the full 3D trajectory of each lipid, which enables resolving leaflet differences and removing the influence projecting lipids onto an imaging plane via the use of a geodesic distance-based analysis of the CG simulations [12]. While the nuances of lipid diffusivity in spatially varying mean and Gaussian curvature are the focus of an upcoming manuscript, here we present the key results of such an analysis. To gain insight into the detailed dynamics of the system, the averaging effect of measurement time was mitigated by extracting the diffusion coefficients for displacements Δ*t* = 1 ns (Fig. S9C) and Δ*t* = 100 ns (Fig. 3G). Both leaflets provided faster diffusion on the bud for phase-separated membranes. However, the use of geodesic distances demonstrated that the experimentally apparent slowing of lipids at 5 and 10 nm from the bud center is indeed due to the 2D projection of the 3D trajectories rather than a reflection of effective local membrane viscosity changes. Furthermore, it was also revealed that the observed increase in lipid mobility correlated well with the mean curvature. Mean curvature explains the non-monotonous diffusion values of the lower leaflet; positive curvature of the membrane neck results in an increase in lipid diffusivity, and the negative curvature of the center of the bud slowed diffusion. For a given lipid type, diffusion rates in the planar membrane region converged to a single value across both leaflets. Hydrodynamics dictates that the effective viscosity in a single phase should be the same for all lipid types [23]. Here, the ratio of diffusion for DLiPC and DPPC lipids was closer to 0.5, which matched the ratio of diffusion coefficients in Martini simulations of planar phase separating systems [40]. This finding is in agreement with the observation that the lipid phases propagated along the flat membrane and spanned the simulation box (Fig. 1G).

## Discussion

This study provides a quantitative analysis of the phase-curvature coupling of SLBs with membrane shapes that are reminiscent of nanoscopic endocytic pits and viral buds. The membrane composition, temperature, and curvature were varied for experimental and simulated membranes. The phase-curvature coupling was revealed by the lateral heterogeneity in both the local density and the diffusion rates of disorder-preferring fluorescent lipids. The curvature influenced the lipids more under conditions that provided a longer tie-line separating the coexisting phases, *i*.*e*., lower temperature and less cholesterol. The effects of curvature were amplified in lipid mixtures even when no phase separation was observed on the planar membrane, which suggest that the curvature was able to induce a phase separation. This is consistent with a curvature-induced phase separation even when the planar membrane has no optically resolvable phase separation and that *T*_mix_ is dependent on the membrane shape. Accordingly, these results suggest that the dynamic curvature and composition of native biological membranes may couple to drive complex membrane processes in membrane signaling, shape changes, and pathophysiology.

### Membrane brightness at curvature sites

The 1:1:2 SLBs phase sorting result indicated that the ratio of *P*_C_*/P*_F_ was greater than one for all values of *P*_F_ (Figs. 2A and S5); the disorder-preferring fluorescent lipid density was higher on curved membranes. The sorting was presumably due to the differences in the phase bending rigidity because the leaflets seemed to maintain phase registration and we do not anticipate that the molecular shape (*e*.*g*., the lipid packing parameter) of the fluorescent lipid was sufficient to induce significant single-molecule, phase-independent sorting [4, 7, 21]. This is further supported by our observation that quasi-one component membranes of POPC were not able to induce a curvature-dependent sorting of fluorescent lipids experimentally or alter the POPC density in CG simulations (Fig. 2E) [49].

### Diffusion reveals the coupling of phases and curvature

The ratio of diffusion on curved versus flat membranes (*D*_C_*/D*_F_) was larger when phase separation was present (Fig. 3). This phenomenon was observed for all values of *P*_F_ for the DiPhyPC, DPPC, and cholesterol membranes, which suggested that the disorder-preferring lipids sorted to curvature even when the curvature was surrounded by the L_d_ phase. This result is consistent with sorting studies where *P*_C_*/P*_F_ values were above 1 for all *P*_F_ (Fig. 2A) and is further supported by CG simulations of DLiPC, DPPC, and cholesterol (Fig. S9). Quasi-one component L_d_ membranes of DiPhyPC displayed no significant difference between *D*_C_ and *D*_F_ whereas mixtures of DiPhyPC, DPPC, and cholesterol displayed faster diffusion at curved surfaces regardless of if phase separation was observed on the planar membrane. We hypothesize that this demonstrates the capability of curvature to induce phase separation that is not present on flat membranes by effectively increasing the local *T*_mix_.

The minimal differences in diffusion observed for membranes of varying phases throughout Fig. 3 are likely due to the intrinsic noise incorporated into the determination of the membrane phase surrounding each membrane bud (Fig. S4). These diffusion studies were practically limited in the number of membrane buds able to be measured, the uncertainty of the phase at 400 nm away from each bud, and the single-lipid diffusion rates measured at each radial distance (Table S1).

### Curvature-seeded phase separation propagates onto the surrounding flat membrane

A subtle but consistent decrease in *D*_*xy*_ was observed on the flat membrane propagating away from the engineered curvature sites (Fig. 3C). This demonstrates that the effective membrane viscosity and the composition of the flat membrane varied with distance from the nanoparticle as if the curvature seeded a phase separation that propagated onto the surrounding flat membrane. This phenomenon was also observed in continuum and CG simulations of phase separation with the L_d_ phase more likely to be present on the flat membrane that is near the curvature versus far from the curvature. (Fig. 2). The simulated L_d_ phase connected across the periodic images along one axis of the simulation box (Fig. 1G). While this indeed matches the phase sorting propagating onto the flat membrane, it is also a trivial consequence of the fixed amount of L_d_ and L_o_ lipids in a finite simulation box.

## Conclusions

Nanoscale membrane curvature was engineered in phase-separated supported lipid bilayers. The sorting and dynamics of lipids were measured to reveal the curvature-induced phase separation and the phase-curvature coupling at physiological length scales. The fluorescence intensity ratio of curvature versus surrounding planar bilayer from diffraction-limited images show the L_d_-preferring lipids sorted to the engineered curvature. The single-lipid diffusion rates were measured versus the lateral distance away from the center of curvature to further support the curvature-induced sorting of disordered lipids. Unlike quasi-one component liquid DiPhyPC or POPC bilayers, SLBs with ternary mixtures of lipids showed a faster diffusion on the curved versus flat membranes. The effects of curvature on lipid diffusion and sorting were more pronounced under conditions in which the phase-separating tie-lines were longer, including lower temperatures and less cholesterol, and when the local surrounding planar membrane was more ordered. These results indicate that the strong preference of disorder-preferring lipids to membranes curved at physiological scales.

## Methods

This manuscript incorporates a diverse collection of experimental and computational methods that are detailed in the Supporting Information. The formation of GUVs, SLBs, and nanoengineered substrates are detailed in sections S1.1 and S1.2. To maintain the large-scale phase separation in the SLBs similar to as observed on the GUVs, required precise temperature control of the sample and substrate was required during SLB formation (S1.3). The diffraction-limited imaging (S1.4) and single-molecule localizations (S1.5) occurred with precise sample temperature control (S1.6). Data analysis included spatial mapping of the lipid phases (S1.7), membrane curvature (S1.8), single-particle tracking (S1.9), and diffusion coefficient calculations (S1.10). Considerations of the effects of the fluorophore (S1.11) and the substrate (S1.12) on the lipid dynamics are presented. Computational approaches include the continuum simulations with explicit phase-curvature coupling (S1.13) and coarse-grained molecular dynamics simulations (S1.14).

## Supporting Information Appendix

This article contains supporting information below. The supplemental movies are available from https://doi.org/10.6084/m9.figshare.c.5928050.

## Acknowledgements

The authors thank Aurelia R. Honerkamp-Smith for valuable discussions. CSC – IT Center of Science is acknowledged for computational resources. X.W. was supported by the Thomas C. Rumble University graduate fellowship, Wayne State University Summer Dissertation Award, and the Richard Barber Interdisciplinary Research Program. M.J. was supported by an Academy of Finland postdoctoral researcher grant (Grant No. 338160) and the Emil Aaltonen Foundation. This material is based upon work supported by the National Science Foundation under Grant No. DMR1652316.

## Author Contributions

X.W., M.J., B.F., and C.V.K. wrote the manuscript. X.W. and C.V.K. designed the experiments and performed the data analysis. X.W. performed the experiments. C.V.K. performed the continuum simulations. M.J. and B.F. performed the CG simulations.

