## Supplemental Materials for "Nanoscale membrane curvature sorts lipid phases and alters lipid diffusion"

### S1 Supplemental Materials and Methods

#### S1.1 GUV formation

The lipids 1,2-diphytanoyl-sn-glycero-3-phosphocholin (DiPhyPC), 1,2-dioleoyl-sn-glycero-3-phosphocholine (DOPC), 1,2-dipalmitoyl-sn-glycero-3-phosphocholine (DPPC), 1-palmitoyl-2-oleoyl-glycero-3-phosphocholine (POPC), and cholesterol (Avanti Polar Lipids) were used without further purification (Fig. 1 in the main text). Fluorescent 1,2-dipalmitoyl-sn-glycero-3-phosphoethanolamine-Texas Red (DPPE-Texas Red, Life Technologies) was included at 0.1 mol% for labeling for all membranes with DiPhyPC or POPC. 1-palmitoyl-2-(dipyrrometheneboron difluoride)undecanoyl-sn-glycero-3-phosphocholine (TopFluor-PC; Avanti Polar lipids) was included at 0.2 mol% in membranes with DOPC. Milli-Q water with a resistivity of 18 mΩ was used in all buffers. All other chemicals were purchased from Sigma Aldrich.

GUVs were created by electroformation, as described previously [6]. Briefly, lipids were combined in chloroform and dried onto conducting indium tin oxide-coated glass plates. A trimmed silicon sheet was added between the plates to form the incubation chamber. The chamber was filled with a 200 mM sucrose solution and subjected to an AC signal with 3 $V_{rms}$ at 10 Hz for 1 hr at 55°C. The GUV solution had 13 mg of lipids per mL after electroformation. GUVs were stored at 55°C and used within 2 days.

#### S1.2 Sample dish preparation

Glass-bottom dishes (MatTek Corp.) were initially rinsed with ethanol, blown dry by a nitrogen stream, then placed in air plasma (Harrick Plasma) for 10 s to create a hydrophilic surface. 20 µL of 5 mM $CaCl_2$ was spun on a glass substrate at 100 rpm. 50-nm radius fluorescent nanoparticles that were excited at 405 nm (Fluoro-Max; Fisher Scientific) were used for generating curvature, and multicolored nanoparticles (TetraSpeck, ThermoFisher Scientific Technologies) were used as stage drift correction fiducial marks. The nanoparticle-containing solutions were evaporated from the coverslips by a hot plate at 35°C for 5 min. The dishes were then chilled to room temperature prior to GUV deposition.

#### S1.3 Supported lipid bilayer formation

SLBs with engineered curvature were created by GUV fusion over nanoparticles on microscopy coverslips, similar to as done previously [4, 9, 10, 21]. However, the new methods for preserving the large-scale phase separation from GUVs in solution to SLB patches on the coverslip were developed here. The GUV solution was placed in the refrigerator at 4°C for 2 minutes to assist the macro-domain formation. 5 µL of the chilled GUV solution was applied to the room temperature glass bottom dishes with nanoparticles. 50 µL of 4°C Milli-Q water was added to encourage the GUVs to sink to the cover glass. The dish was chilled to 4°C for 15 min before gentle rinsing with 5 mL of 4°C, 200 mM sucrose.

#### S1.4 Imaging procedure

The optical setup included an inverted IX83 microscope with a 100×, 1.49 NA objective (Olympus), a 2× emission path magnification (OptoSplit, Cairn Research), and an iXon

897-Ultra EMCCD camera (Andor Technology), as described previously [9–11, 21]. A Hg lamp with an excitation filter (BrightLine single-band filters, Semrock) provided illumination for diffraction-limited images (*i.e.*, Fig. S1). CUBE diode lasers with wavelengths of 405 and 488 nm (Coherent) and a 561 nm Sapphire laser (Coherent) were used for single-molecule localization microscopy. The excitation light passed through a clean-up filter (zet405/488/561/647x, Chroma Technology), encountered a quad-band dichroic mirror (zt405/488/561/647rpc, Chroma Technology), and reflected into the objective. The emission was isolated via emission filters (BrightLine single-band filters, Semrock) and a 4-band notch filter (zet405/488/561/640m, Chroma Technology). The SOLIS imaging software (Andor Technology) was used to acquire images with 128 pixels x 128 pixels region of interest in the kinetic mode and an EM gain of 150. The images for single-molecule localization were acquired at 537 Hz. Typically, 20,000 frames were combined to create reconstructed, super-resolution images or analyzed for single-particle tracking.

The degree of lipid sorting with curvature was quantified by measuring the fluorescence intensity difference between the engineered curvature and the surrounding planar membrane via diffraction-limited epifluorescence imaging. The membrane intensity in diffraction-limited images represents the fluorophore concentration with 200 nm resolution. Diffraction-limited imaging requires two orders of magnitude less illumination intensity than super-resolution imaging. Also, our super-resolution methods show only a few fluorophores per frame and required >20,000 frames to study the diffusion or to reconstruct images. Each curvature site of a POPC membrane provided between 1–30× higher density of DPPE-Texas Red localizations than the planar bilayer, which is of significantly greater variation between buds than observed via diffraction-limited imaging, 1 to 2.5× (Fig. 1A in the main text). Therefore diffraction-limited images were used for phase-sorting studies.

#### S1.5 Single-fluorophore localization

The movies with optically isolated fluorescent lipids were analyzed by the Fiji plug-in ThunderSTORM, which fit every single-fluorophore image with a 2D Gaussian function to export its location, intensity, fit width, and fit certainty [16]. Only the localizations with intensity >100 photons, Gaussian fit width >15 nm, and location uncertainty <45 nm were kept for further analysis. The ThunderSTORM-reported fluorophore uncertainty ($\sigma_r$) was $24 \pm 1$ nm. The stage drift during imaging was corrected by analyzing the motion of the multicolored nanoparticles through built-in ThunderSTORM fiducial tracking, which was separate from the nanoparticles used to engineering membrane curvature.

Super-resolution images reconstructed from single-fluorophore localizations contained aggregates of DPPE-Texas Red in the DiPhyPC-containing membranes. The clusters of these dense, aggregate localizations were $63 \pm 17$ nm radius and showed confinement in lipid diffusion. DiPhyPC contains phytanoyl acyl tails that are highly disordered and have reduce photo-oxidization compared to unsaturated lipids. Photo-stability is especially appreciated in experiments that rely on phase separations that are sensitive to the membrane composition.

Aggregates were identified and removed by custom algorithms that were studied extensively previously [20]. Aggregates were removed via binning all localizations into voxels of space and acquisition time and culling based on a threshold localization rate. When the number of localizations in a voxel exceeded the localization density threshold ($\rho_{\text{th}}$), that region of the sample was deemed to be an aggregate, and all localizations within that $XY$-region were excluded from subsequent analysis. Further, all localizations associated with trajectories that lasted more than 32 steps were culled. The value of $\rho_{\text{th}}$ was varied such that between 0 and 80 percentiles of the total localizations were culled. For each $\rho_{\text{th}}$, a simulated membrane was generated with an evenly distributed localizations except the removed region. A spatial correlation function was calculated for both the experimental and simulated membrane and the remaining localizations according to

$$g(r) = \frac{\left\langle \text{FFT}^{-1}\left(|\text{FFT}(I(\mathbf{r}))|^2\right)\right\rangle_\theta}{\rho^2}. \tag{S1}$$

$I(r)$ represents the two-dimensional localization histogram, and $\rho$ is the average localization density. The average number of localizations per cluster can be calculated by comparing the spatial correlation of the two.

$$N = \int \left(\frac{g_{\text{exp}}}{g_{\text{sim}}} - 1\right) r\text{d}r. \tag{S2}$$

$N$ was calculated for all localization data sets with varying $\rho_{\text{th}}$. The minimum $\rho_{\text{th}}$ for which $N \leq 3$ was used for the diffusion studies.

#### S1.6 Membrane temperature control

A Peltier temperature control dish holder (QE-1HC, Warner Instruments) was used with a custom, insulated dish cover with a thermocouple mount. Once the dish cover is placed on the sample dish, the thermocouple was <0.5 mm above the center of the glass coverslip. The temperature was actively controlled via a custom LabVIEW program. Also, the dishes were never heated above 45°C to protect the microscope objective. When changing temperature, the temperature was changed at 0.5°C/min and remained at the set temperature for 30 min before imaging. The Peltier dish holder was initially set to 10°C before inserting a membrane sample. The Peltier temperature was set to be at 10°C, 30°C, and 45°C resulted in the thermocouple measuring a temperature of $17 \pm 3$°C, $27 \pm 1$°C, and $37 \pm 1$°C, respectively.

#### S1.7 Lipid phase identification

The 2D projected image of a homogeneous membrane yields an increased brightness at the location of membrane curvature compared to the surrounding planar membrane due to the increase in the projected membrane area. Coincident nanoparticles confirmed the membrane curvature in the appropriate color channel. A single 2D Gaussian fit was used to find the center of membrane curvature in the membrane color channel.

The lipid phase was quantified by the partition coefficient of the fluorescent lipids. Both DPPE-Texas Red and TopFluor-PC are $\text{L}_\text{d}$ preferring lipids [12, 17], and the higher local concentration was interpreted as a greater degree of disorder in the acyl tails. The intensity of the curved membrane ($I_\text{C}$) is defined as the average intensity of the nine pixels within 170 nm from the center of curvature, and the intensity surrounding the curvature ($I_\text{F}$) is defined as the average intensity of the pixels $400 \pm 56$ nm away from curvature from diffraction-limited images (Fig. S5). The increased membrane brightness due to the membrane topography was determined from homogeneous POPC membranes and analytical models of the membrane shape, which provide the expected membrane curvature intensity increase due to the increased membrane area in the $XY$-projection, $I_\text{C}^\text{POPC}/I_\text{F}^\text{POPC} = 1.23$. The average membrane intensity ($< I >$) is defined as the midpoint of the mean $\text{L}_\text{d}$ phase intensity ($I_\text{d}$) and mean $\text{L}_\text{o}$ phase intensity ($I_\text{o}$) from each planar SLB patch; $< I >= (I_\text{d} + I_\text{o})/2$. This definition of $< I >$ accounts for individual fields of view that happened to contain more of one phase than the other. When no phase separation was present on the flat membrane, $< I >= I_F$. The lipid phase on the flat membrane surrounding the curvature ($P_F$) and the phase of the curvature ($P_C$) according to

$$P_F = I_F / \langle I \rangle, \tag{S3}$$

$$P_C = \frac{I_C}{\langle I \rangle} \frac{I_F^{POPC}}{I_C^{POPC}}. \tag{S4}$$

This method of determining the lipid phase from fluorescence images accounts for sample-to-sample variation in fluorophore concentrations and temporal changes caused by fluorescence bleaching. The phase surrounding the membrane curvature was determined $r = 400$ nm (Fig. S4) with $P_F < 0.9$ indicating the $L_o$ phase and $P_F > 1.1$ indicating the $L_d$ phase. Homogeneous, multicomponent membrane at a temperature above $T_m$ is recognized as a single fluid-phase membrane. This classification was used for sorting the curvature sites based on their surrounding lipid phase for diffusion analysis (Fig. 3 in the main text). The $P_F$ values varied depending on $r$, which resulted in an uncertainty of the SLB phase and yielded no difference in the DPPE-Texas Red diffusion at $r = 600$ nm. A more precise analysis of the effects of $P_F$ is shown in with diffraction-limited imaging when the classification of the phases was not necessary, and the value of $P_F$ surrounding each nanoparticle could be directly reported (*i.e.*, Fig. 2 in the main text). However, the diffusion analysis required combining the data from multiple nanoparticles to yield statistically meaningful results. A more robust analysis of the phase-dependent diffusion of fluorescent lipids on planar SLBs was provided previously [20].

#### S1.8 Super-resolved curvature identification and aggregate removing method

Reconstructed super-resolution images of single lipids revealed increased localization density at the sites of membrane curvature. Curvature was confirmed by the colocalization of fluorescent nanoparticles and lipid accumulations in the complementary color channels. The local density increase in single-lipid localizations was fit to a 2D Gaussian function to find the exact curvature center for the super-resolution images and single-particle tracking analysis.

DiPhyPC-containing SLBs yield aggregates of DPPE-Texas Red [20]. Aggregates were apparent via a high fluorophore localization density in the absence of nanoparticles. Comparisons between membrane curvature events and fluorophore aggregates were made via the size of the cluster of localizations, the local single-lipid diffusion rate, localization density, localization rate, and the proximity to the nanoparticles. Aggregates were removed via localization density and long trajectory culling, as described in the Supplemental Material and previously [20]. Briefly, all localizations that were not associated with membrane curvature were grouped based on their 2D location and acquisition time into 3D voxels. When the number of localizations in a voxel exceeded the localization density threshold ($\rho_{th}$), that region of the sample was deemed to be an aggregate and all localizations within that $XY$-region were excluded from subsequent analysis. Further, all localizations associated with trajectories that lasted more than 32 steps were culled.

#### S1.9 Single-particle tracking

The single-lipid localizations were linked via u-track with a maximum step length of 400 nm [6]. The average trajectory length of the aggregate-removed localizations was $6 \pm 4$ steps. Only the single-lipid steps between adjacent frames ($\nu$) were used in our analysis while being versus their lateral radial distance away from the center of curvature ($r$) and fit to a 2D Rayleigh distribution ($R$) [9, 20],

$$R(\nu) = \frac{\nu}{2D_{\text{fit}}\Delta t} \exp\left(-\frac{\nu^2}{4D_{\text{fit}}\Delta t}\right). \tag{S5}$$

Fitting of Eq. S5 incorporates the time between sequential frames ($\Delta t$) and yields a fit diffusion coefficient ($D_{\text{fit}}$). $D_{\text{fit}}$ was corrected for imaging blur created by the single-frame exposure time ($t_{\text{exp}}$) and localization uncertainty ($\sigma_r$) [2] according to

$$D_{xy} = \frac{D_{\text{fit}} - \frac{\sigma_r^2}{2\Delta t}}{1 - \frac{t_{\text{exp}}}{3\Delta t}}. \tag{S6}$$

When Eq. S6 was applied to CG simulations with a $\sigma_r > 0$ applied to mimic the experimental conditions, $t_{\text{exp}} = 0$ because the instantaneous lipid COM was calculated without blurring over an exposure time.

#### S1.10 Monte Carlo fitting of diffusion

The measured lateral diffusion rates ($D_{xy}$) vs. radial distance from the nanoparticle center ($r$) includes contributions from both the effects of local curvature on the lipid mobility and the 3D membrane shape projected into the $XY$-imaging plane. To extract the separate contribution of these two effects, Monte Carlo simulations were performed [9, 21]. Individual lipids were simulated diffusing on a curved surface that estimates the shape of the SLB draped over the nanoparticle on cover glass S6. First, available locations for the lipid on the membrane were created at a density of 4/nm$^2$. Second, a single-lipid trajectory was calculated by allowing the lipid to randomly step to one of the $110 \pm 10$ points within 3 nm of its current location with an average step length of 2 nm. These step lengths were significantly smaller than the surface's radius of curvature. A duration of time was associated to each single lipid step ($\Delta t_1$) to set a local effective diffusion coefficient. For example, when $\Delta t_1 = 0.45$ µs, then a local diffusion coefficient in the membrane of 2.5 µm$^2$/s was simulated. This local diffusion coefficient represents the in-plane diffusion for the membrane, regardless of the membrane's local orientation relative to the $XY$-plane. Varying curvature-induced changes to lipid diffusion were simulated by mimicking a different local D, *i.e.* spatially dependent $Deltat_1$ values. The simplifying assumption was made to reduce fitting parameters that all lipid diffusion on the planar membrane had one diffusion coefficient and all membrane with curvature had a second diffusion coefficient.

To connect these Monte Carlo simulations to the experimental data, trajectories were calculated until the cumulative time of 1.86 ms had elapsed, which is the frame rate of our experimental single-molecule localization and SPT. If a constant $\Delta t_1 = 0.45$ µs was used, then a single-lipid trajectory included 413 steps. When $Deltat_1$ varied during the lipid trajectory, the number of simulated steps in a single trajectory varied. The random trajectories starting and ending locations on the membrane were saved and analyzed as if they were sequential frames in a SPT experiment. Additionally, a 20 nm standard deviation Gaussian localization imprecision was applied. The single-lipid step lengths were projected into the $XY$-plane and analyzed vs. lateral radial distance from the curvature center across the sample were analyzed just as was done for the experimental data with $t_{exp} = 0$. A matrix of Monte Carlo simulations were performed with varying $\Delta t_1$ for the planar membrane and $\Delta t_1$ for the curved membrane. This matrix of Monte Carlo results were fit to the experimentally acquired $D_{xy}$ vs. $r$ to extract the effective diffusion coefficient on the flat and planar membranes. The extensive, dense sampling of quasi-single-component membranes revealed a consistent geometrical effect of $XY$-projection on the observed $D_{xy}$ vs. $r$. The simulations show that $D_{xy}(r \leq 50$ nm) underestimates the in-membrane diffusion by a factor of $0.53 \pm 0.10$ due to the membrane tilt relative to the $XY$-plane for membranes wrapping the 50-nm radius nanoparticles. The single-fluorophore localizations could not feasibly be acquired a densely for the curved, phase-separated membranes presented in this manuscript. Accordingly, the diffusion of the curved ($D_{\mathrm{C}}$) and flat ($D_{\mathrm{F}}$) membranes is approximated by

$$D_{\mathrm{C}} = D_{xy}(r \leq 50 \text{ nm})/0.53, \tag{S7}$$

$$D_{\mathrm{F}} = D_{xy}(400 \text{ nm} \leq r \leq 600 \text{ nm}). \tag{S8}$$

#### S1.11 Effects of the fluorescent lipid

The lipid diffusion with nanoscale curvature varies with the fluorescence labeling strategy. For example, DPPE-Texas Red lipid diffusion in POPC bilayer in a curved membrane displayed a diffusion rate 40% of that of the flat membrane, $D_{\mathrm{C}}/D_{\mathrm{F}} = 0.4 \pm 0.1$. However, TopFluor-PC showed no difference in diffusion with membrane bending, $D_{\mathrm{C}}/D_{\mathrm{F}} = 1.0 \pm 0.2$ [21]. DPPE-Texas Red was primarily used rather than TopFluor-PC throughout this manuscript because it was brighter, it could be used at lower concentrations, and it required less intense fluorescence illumination power. Moreover, tail-labeled fluorescent lipids perturb membrane more than head group-labeled fluorescent lipids as measured by transient binding versus free diffusion, which indicates the level of trapping interactions [15].

#### S1.12 Effects of the substrate on lipid diffusion

The diffusion of individual lipids and phase domains in SLBs are affected by the substrate's close proximity to the bilayer's lower leaflet. This substrate proximity slows the diffusion of single lipids in SLB to 10–50% of the observed diffusion in vesicles or black lipid membranes [13]. Similarly, the diffusion in the bilayer's top leaflet, distal from the substrate, is 1.1–1.3× faster than the single-lipid diffusion in the bottom, proximal leaflet [21]. Whereas the substrate roughness and surface chemistry can greatly affect lipid diffusion, we have examined the SPT of fluorescent lipids in the leaflets separately to confirm that the nanoparticle vs. coverslip surface properties do not contribute to the $D_{\mathrm{C}}/D_{\mathrm{F}}$ values reported in this manuscript. [21].

#### S1.13 Continuum simulations with explicit phase-curvature coupling

Lipid phase separation and phases-curvature coupling was modeled with a Hamiltonian consisting of three parts: membrane height fluctuations via the Helfrich form; local compositional fluctuations via a fourth-order Landau expansion; and curvature-composition coupling [18],

$$H = \int \left[\kappa \left(\nabla^2 h\right)^2 + \sigma \left(\nabla h\right)^2 + A\phi^2 + B\left(\nabla \phi\right)^2 + C\phi^4 - \gamma\phi\left(\nabla^2 h\right)\right] \mathrm{d}x\mathrm{d}y. \tag{S9}$$

The Hamiltonian is dependent on the membrane height ($h$), bending rigidity ($\kappa$), the surface tension ($\sigma$), phase ($\phi$), Landau phase constants ($A$, $B$, and $C$), and phase-curvature coupling strength ($\gamma$). Our simulation examined the distribution of lipid phases on the membrane of fixed curvature, similar to as done experimentally

(Fig. S6). Both $\kappa$ and $\sigma$ were set to zero because the membrane shape was held constant and $h$ was unchanging during the simulation; any constant values of $\kappa$ and $\sigma$ would cancel when calculating changes in $H$. The phase-dependent bending rigidity is incorporated within $\gamma$ as one of the possible physical explanations of a phase-curvature coupling and a non-zero value for $\gamma$. The Landau phase constants $A$, $B$, and $C$ enable modeling the phase separation away from the critical point where first-order phase transitions are expected. We set $A = -1$, $B = 0.7$, and $C = 4$; a minimum $H$ was observed on flat membranes (*i.e.*, $\nabla^2 h = 0$) at $\phi = \pm 2 - 3/2$ and the phases were well separated with minimal contact length (*i.e.*, small $(\Delta\phi)^2$). Positive values of $\gamma$ sorted positive values of $\phi$ to the curvature, which we labeled as $\mathrm{L_d}$. A time step of our simulation included minimizing $H$ by small changes to $\phi$ equal to $-0.05 \times \partial H/\partial\phi$ on location of the membrane. The membrane was divided into a continuum of 9 $\mathrm{nm}^2$ cells. Thermal fluctuations were modeled as the addition of a normal distribution of random perturbations to $\phi$. The standard deviation of this noise distribution ($\sigma$) corresponds to temperature ($T$), which was calibrated by qualitative matching of the experimental observations; flat bilayers displayed no stable phase separation at $\sigma = 0.08$, but large phase domains were present at $\sigma = 0.07$; we observed a qualitative similarities between these two $\sigma$ values and mixed lipid experiments conducted at $T = 15$°C and 35°C, respectively. Additionally, the perimeter of each simulation ($r \geq 150$ nm) was rescaled each time step to have a $< \phi >= 0$ to prevent the simulation from becoming a single large domain of one phase at low temperatures. Movies of the phase dynamics and a static image of the average phase separation on flat and curved membranes are provided in the Supporting Information with varying $T$ and $\gamma$ (Movie S1 and Fig. S7).

#### S1.14 Coarse-grained molecular dynamics simulations

Coarse-grained (CG) Martini [14] simulations were performed with the GROMACS 2020 package [1] with full hydration, and 10% of the solvent was modeled as the antifreeze particles to prevent the well-known crystallization of the solvent in the Martini force field [14]. The curved membranes were generated by the BUMPy script [3] and maintained by dummy particles that repelled the acyl chains. The dummy participles maintained a hemispherical bud of 5 nm radius connected to a planar bilayer *via* a neck of minimum 5 nm radius of curvature. The two compositions simulated were (1) pure POPC (4497 lipids and 148605 water beads) and (2) a phase separated mixture of 2175 disorder-preferring 1,2-dilinoleoyl-sn-glycero-3-phosphocholine (DLiPC), 2216 order-preferring DPPC, and 1524 cholesterol molecules with 158227 water beads. The systems were simulated at 37°C for 20 µs, of which the final 10 µs were included in the analyses. Five replicates were performed for the ternary lipid mixture. The molecular trajectories are freely available for download [7, 8]. Analysis of the CG results included tracking each lipid type while mimicking the analysis procedure performed on the experimental data, including adding a radial localization imprecision ($\sigma_r$) designed to mimic the experimental localization uncertainty. Because the bud radius in experimental systems is 10× larger than that in CG simulations, the most interesting comparisons are that with a $\Delta t$ that was 100× longer (*i.e.*, 2 ms vs. 20 µs) and $\sigma_r$ that was 10× larger (*i.e.*, 20 nm vs. 2 nm) in experiments than simulations. The analysis was also repeated using a novel, geodesic distance-based algorithm to compute Mean Square Displacements instead of the 3D or projected 2D displacements. In the method, the lipids were separated into leaflets. Due to its high rate of flip-flops, cholesterol molecules were excluded from the analysis. Triangular mesh surfaces were created based on the individual leaflets with triangle sizes smaller than 8 Å, and the last 10 µs of lipid displacements were mapped onto the surface. Finally, the corresponding geodesic distances were computed. For further details of the method, see reference [5].

#### S2 Supplemental Figures

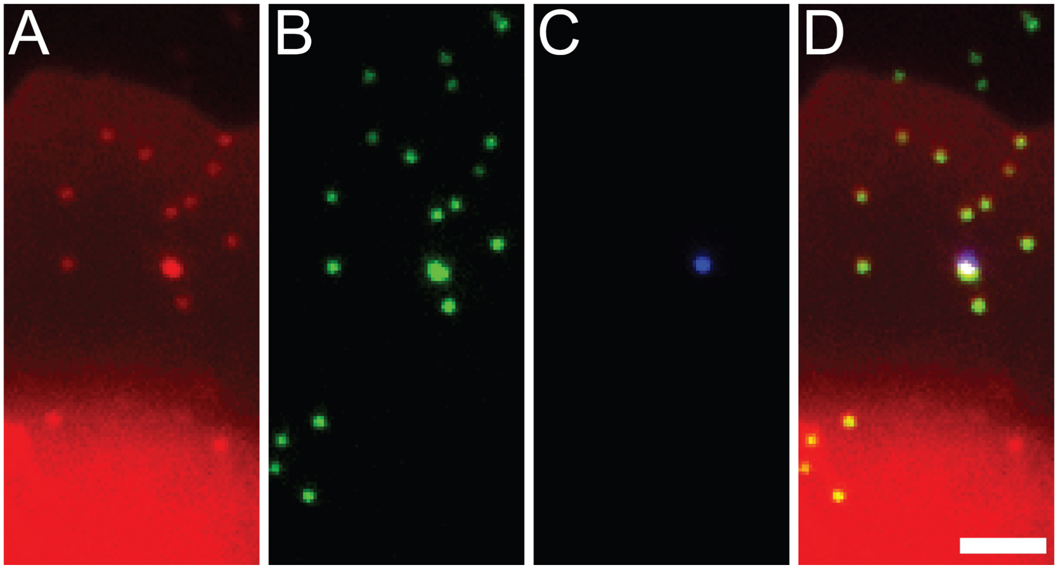


Figure S1: Phase-separated SLBs were created with nanoscale membrane curvature. Here, a 1:1:2 molar ratio of DiPhyPC:DPPC:cholesterol and 0.1 mol% DPPE-Texas Red is shown over 50-nm radius nanoparticles. Diffraction-limited fluorescence images include those of (A) DPPE-Texas Red, (B) the fluorescent nanoparticles, and (C) the multicolored nanoparticle fiducial markers, and (D) their merge. (A) The phase-separated SLB contained a micron-scale $L_d$ domain (bright, bottom) and $L_o$ domain (dim, middle) while surrounded by bare coverslip (black, top). Nanoscale puncta represent an increased membrane area and a sorting of the $L_d$ phase where membrane curvature is present, such as the outlined region of (A) that is shown as a zoom-in (Fig. 1C). Scale bar, 2 µm.

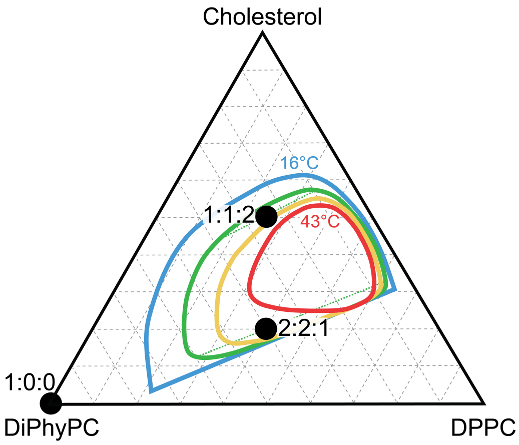


Figure S2: Lipid bilayers composed of mixtures of DiPhyPC, DPPC, and cholesterol displayed two coexisting liquid phases at select compositions. The mixtures of DiPhyPC:DPPC:cholesterol used in this study are shown (*black dots*). The 16°C (*blue*) and 43°C (*red*) phase boundaries were measured previously via fluorescence microscopy of GUVs [19]. The boundary of liquid phase coexistence at 25°C (*green*) and 34°C (*yellow*) were approximated. At 25°C, the tie-line (*green dotted*) is longer for the composition of 2:2:1 than 1:1:2, indicating a greater difference between the coexisting liquid phases for 2:2:1 vs. 1:1:2. This figure is adapted from our prior publication, Ref. 20

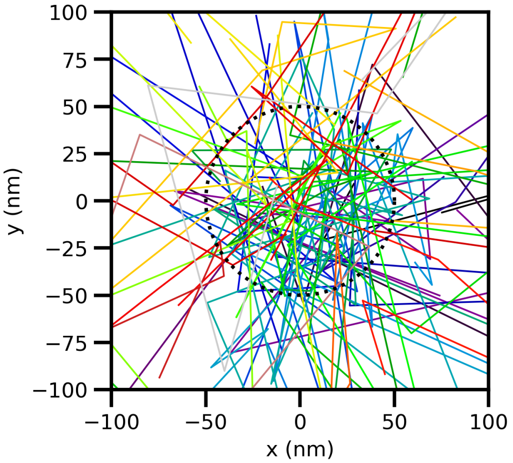


Figure S3: The membrane draped over the nanoparticles upon the coverslip yields membrane buds connected to the planar SLB. This is demonstrated by fluorescence recovery after photobleaching [4,10] and single-lipid trajectories. Single-lipid trajectories, such as those shown here, traverse between the planar SLB and the membrane over the nanoparticle.

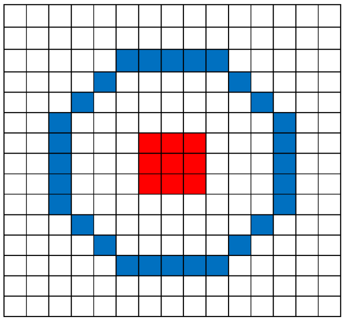

Figure S4: Diffraction-limited images were used to assess the sorting of disorder-preferring fluorescent lipids to the curved membranes. The curvature center was identified by the colocalization of the fluorescent nanoparticle and the increase in fluorescent lipids in the $XY$-projected image. Grid shown here was overlaid with the projected image of fluorescent lipids to measure the brightness of the curvature and the brightness of the surrounding, flat membrane. The center 9 pixels (*red*) were deemed to be equal to the curved membrane and the pixels that were closest to 400 nm from the nanoparticle center were deemed equal to the surrounding flat membrane (*blue*). Each pixel maps to 80 nm × 80 nm of the sample.

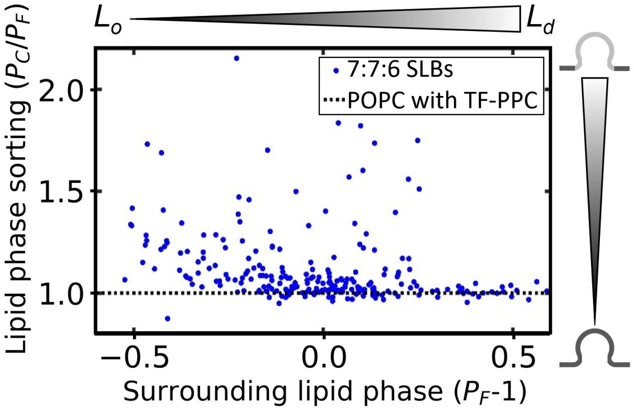


Figure S5: Curvature-induced lipid phase sorting was confirmed in SLBs made with a 7:7:6 molar ratio of DOPC:DPPC:cholesterol and labeled with the fluorescent lipid TopFluor-PPC, which concentrates in disordered lipid phases. $P_C/P_F$ was greater than 1, indicating that disorder-preferring lipids sorted to the curved membranes. Similar to Fig. 2A in the main text, the curvature-induced sorting was most significant for curvature regions that were immediately surrounded by order-preferring lipids; $P_C/P_F$ decreased with increasing $P_F$.

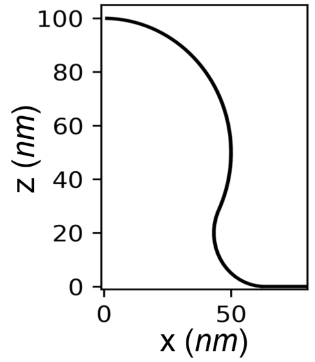


Figure S6: Continuum simulations of membrane phase and Monte Carlo simulations of lipid diffusion occurred on the azymuthially symmetric membrane shape shown here. The spherical bud top includes a 50-nm radius of curvature. The bud neck smoothly connects the spherical top to the planar surroundings with a minimum 20-nm radius of curvature. This shape is also the best estimate of the experimental membrane shape over the 50-nm radius nanoparticles.

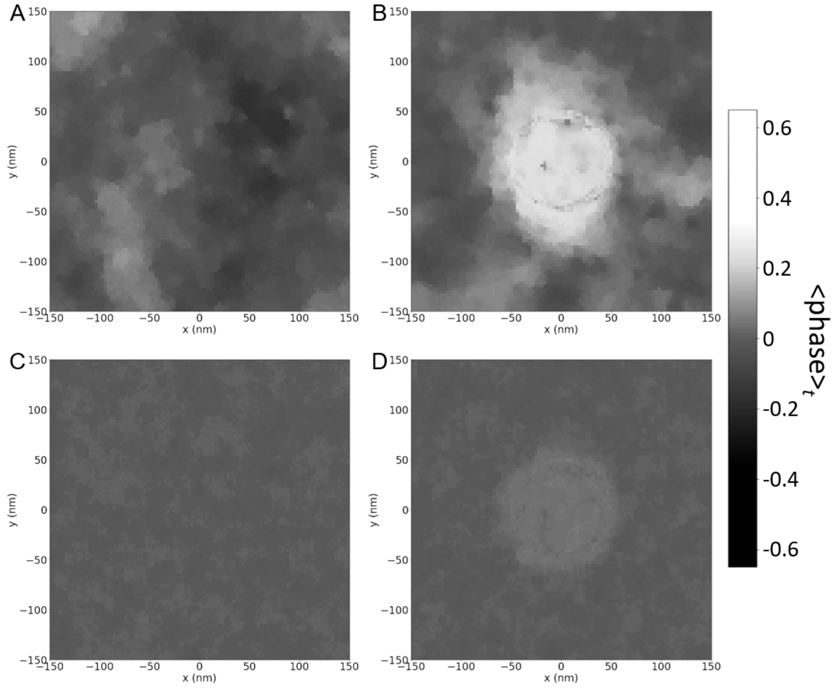


Figure S7: The continuum simulations include phase fluctuations and sorting of the disordered phase to the center, curved membrane with a balance of the Landau phase constants, phase-curvature coupling, and thermal fluctuations. Shown here is top-view of the time-averaged membrane phase ($\phi$), as indicated by the gray-scale. These simulations included (A, B) cold ($\sigma = 0.07$) and (C, D) hot ($\sigma = 0.08$) membranes that are (A, C) flat or (B, D) with a 50-nm radius membrane bud with a phase-curvature coupling ($\gamma$) of 0.1. These data are the time-averaged data that is also shown in Fig. 2 in the main text and Movie S1.

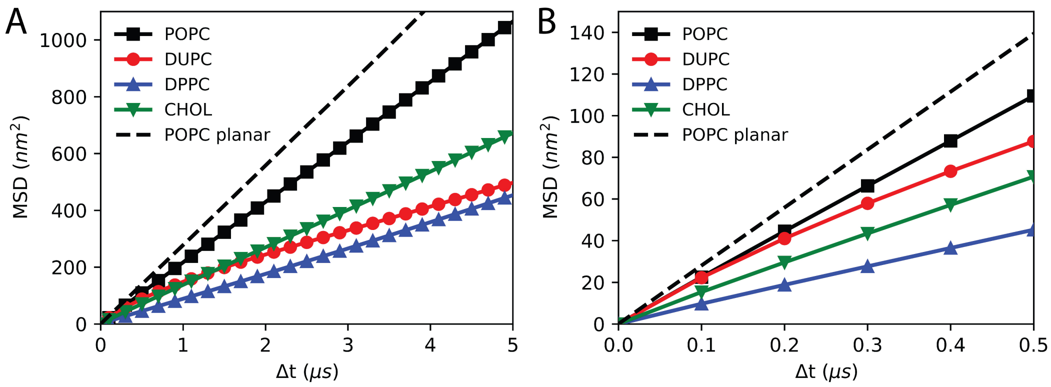


Figure S8: A conventional mean squared displacement (MSD) vs. $\Delta t$ analysis of the CG lipid diffusion on the membrane bud, unless otherwise noted. Simulations with periodic boundary conditions were unwrapped to ensure all lipid trajectories were intact before linking and analysis. The presence of the membrane curvature resulted in a non-Brownian (*i.e.* non-linear) MSD vs. $\Delta t$. The averaging of many long trajectories into each of these data sets resulted in the curved and planar lipid mobility incorporated into these curves such that extracting the lipid diffusion on the curved vs. flat membrane was impossible. (B) shows the zoom of of (A) at short $\Delta t$ values.

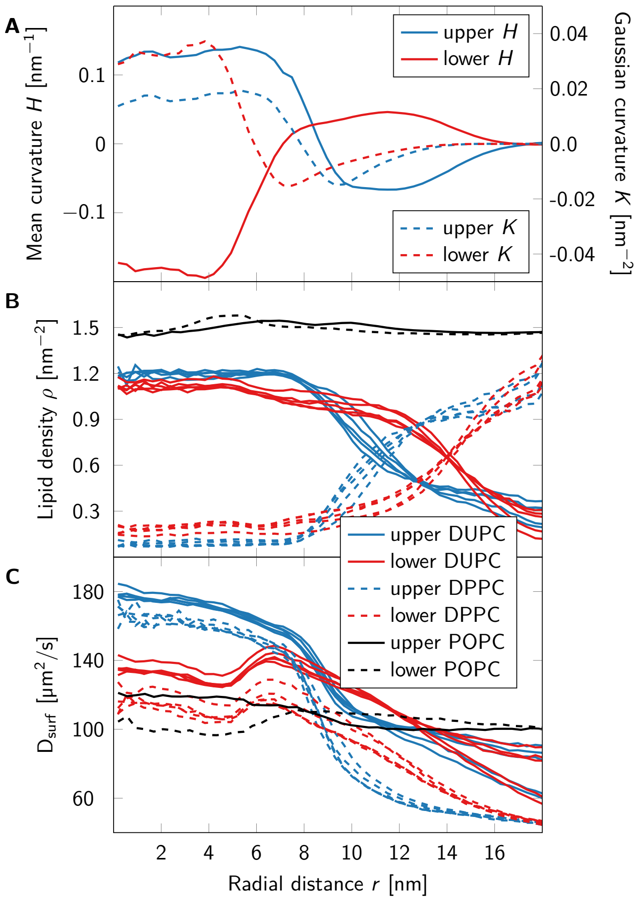


Figure S9: Geodesic distance-based analysis of lipid displacements in the CG simulations of DLiPC, DPPC, and cholesterol. (A) Leaflet-wise mean ($H$) and Gaussian ($K$) curvatures of the simulated membrane bud. (B) The on-surface densities of POPC, DPPC, and DLiPC lipids, as observed in the individual leaflets, allow the determination of membrane composition in regions of various curvature (C) The diffusion coefficient through the modeled membrane surface ($D_{\mathrm{surf}}$) was computed from displacements at $t$ = 1 ns highlights the differences between lipid diffusivity in the upper and lower membrane leaflets. Budding membranes also possess mean curvature whose sign is conventionally taken as positive if the membrane bends locally away from the outer leaflet. Because this definition is ill-defined for non-closed surfaces, here, a leaflet-wise approach was adopted: the mean curvature of a leaflet was positive when the membrane curved away from the lipid head groups. Using this definition, the mean curvature correlated with the Gaussian curvature in absolute value, but it had different signs for the upper and lower leaflets.

#### S3 Supplemental Movies

The supplemental movies are available from
https://doi.org/10.6084/m9.figshare.c.5928050.

##### Movie S1

The phase dynamics at (A, B) cold ($\sigma = 0.07$) and (C, D) hot ($\sigma = 0.08$) on (A, C) flat and (B, D) curved membranes as governed by Eq. S9. Snapshots of (B, D) are shown in Fig. 1E, F. The the static curved membrane shape in (B, D) is shown in Fig. S6.

##### Movie S2

Trajectories of a mixed DLiPC, DPPC, and cholesterol within a fixed membrane topography subjected to the Martini force field. The lipids and bead type are shown in Fig. 1 in the main text. Each frame here corresponds to a 100 ns step of simulation time.

#### S4 Supplemental Tables

Table S1: Diffusion results for varying membrane compositions, temperatures, and phases. The $D_{\mathrm{F}}$ uncertainties are from the standard deviation $D_{xy}$ values for $r$ between 400 and 600 nm. The $D_{\mathrm{C}}/D_{\mathrm{F}}$ and uncertainties are from the propagated error of $D_{\mathrm{C}}$ and $D_{\mathrm{F}}$ fitting. The listed membrane compositions are the molar ratios of DiPhyPC:DPPC:Cholesterol, unless otherwise stated.

| Composition | $T$ (°C) | Phase | Num. of samples | $D_{\mathrm{F}}$ (μm$^2$/s) | $D_{\mathrm{C}}/D_{\mathrm{F}}$ |
|---|---|---|---|---|---|
| POPC | 25 | $L_d$ | 12 | $2.6 \pm 0.03$ | $0.4 \pm 0.1$ |
| DiPhyPC (1:0:0) | 17 | $L_d$ | 19 | $1.03 \pm 0.03$ | $1.2 \pm 0.4$ |
| DiPhyPC (1:0:0) | 36 | $L_d$ | 18 | $1.75 \pm 0.03$ | $0.7 \pm 0.4$ |
| 1:1:2 | 19 | $L_o$ | 8 | $0.78 \pm 0.04$ | $2.3 \pm 1.7$ |
| 1:1:2 | 19 | $L_d$ | 8 | $0.74 \pm 0.13$ | $2.1 \pm 1.4$ |
| 1:1:2 | 27 | $L_o$ | 14 | $1.27 \pm 0.09$ | $1.5 \pm 0.5$ |
| 1:1:2 | 27 | $L_d$ | 8 | $1.28 \pm 0.07$ | $1.3 \pm 0.4$ |
| 1:1:2 | 37 | $L_o$ | 11 | $1.44 \pm 0.09$ | $1.4 \pm 0.5$ |
| 2:2:1 | 17 | $L_o$ | 11 | $0.67 \pm 0.04$ | $1.7 \pm 0.7$ |
| 2:2:1 | 17 | $L_d$ | 14 | $0.57 \pm 0.06$ | $2.3 \pm 1.0$ |
| 2:2:1 | 27 | $L_o$ | 10 | $1.19 \pm 0.06$ | $1.1 \pm 0.7$ |
| 2:2:1 | 27 | $L_d$ | 16 | $1.37 \pm 0.03$ | $1.5 \pm 0.6$ |

[3] K. J. Boyd and E. R. May. Bumpy: A model-independent tool for constructing lipid bilayers of varying curvature and composition. *Journal of Chemical Theory*

*and Computation*, 14(12):6642–6652, 12 2018. PMID: 30431272 PMCID: PMC6545900.

[4] P. P. Cheney, A. W. Weisgerber, A. M. Feuerbach, and M. K. Knowles. Single lipid molecule dynamics on supported lipid bilayers with membrane curvature. *Membranes*, 7(1), 3 2017. PMID: 28294967 PMCID: PMC5371976.

[5] B. Fábián and M. Javanainen. Curd: A tool for diffusion analyses on curved membranes. *ChemRxiv*, Oct 2021.
